# Evaluation of fluorescent proteins for compatibility with STED microscopy systems using two-color spectroscopies

**DOI:** 10.64898/2026.05.11.724171

**Authors:** Keisuke Sato, Daisuke Okada, Ayana Sugizaki, Tatsuo Nakagawa, Hiroshi Kumagai, Yoshinori Iketaki, Sumio Terada

**Author notes:** Corresponding authors (Y.I.), (S.T).

## Abstract

Stimulated emission depletion (STED) microscopy is a super-resolution fluorescence imaging technique that achieves high spatial and temporal resolution by exploiting stimulated emission to induce fluorescence depletion (FD) and is expected to have substantial utility for imaging applications using fluorescent proteins. However, the compatibility of fluorescent proteins with STED microscopy systems has been understood primarily through empirical observations, and there is no established methodology for the rational selection of fluorescent proteins for STED microscopy. In this study, we systematically evaluated the compatibility of commonly used fluorescent proteins with STED microscopy systems by measuring FD properties using transient absorption spectroscopy and fluorescence dip spectroscopy, both of which are classified as two-color spectroscopy (TCS). Fluorescent proteins identified as compatible with the STED microscopy system based on the TCS measurements were employed for three-dimensional STED imaging of cellular samples expressing each protein. In all samples, three-dimensional spatial resolution was improved relative to confocal laser microscopy, with particularly marked improvements in z-axis resolution. These findings demonstrate that measurements of FD properties via TCS provide a robust approach for evaluating the compatibility of fluorescent proteins with the STED microscopy system and for selecting suitable fluorescent proteins for STED imaging.

## Introduction

Fluorescence imaging enables the specific visualization of targets of interest within biological samples, such as tissues and cells, by labeling them with fluorescent dyes or fluorescent proteins. Recent advancements in information processing and optics have significantly progressed fluorescence imaging. Traditionally, the spatial resolution of fluorescence microscopy was believed to be limited by the diffraction limit, approximately 200 nm at most. However, since the beginning of this century, various super-resolution fluorescence microscopy technologies have been established that can provide spatial resolution surpassing this limit^1–3^. Among them, stimulated emission depletion microscopy (STED) is distinctive in that it is fundamentally based on spectroscopy^4^. In STED, the first laser (pump beam) excites fluorescent molecules from the ground state (S_0_) to the excited state (S_1_), where fluorescence light can be emitted. Next, the second laser (erase beam) quenches molecules in the S_1_ state via stimulated emission, thereby preventing fluorescence emission (fluorescence depletion: FD)^5,6^. In STED, the erase beam is focused into a doughnut-shaped spot together with the usual Gaussian-shaped pump beam^7,8^. When these beams are focused onto a sample labeled with fluorescent molecules, FD takes place in the overlapping area of the beams, thereby generating a fluorescence spot smaller than the diffraction limit in the immediate vicinity of the central point with the maximum fluorescence intensity. By scanning the sample with this spot, fluorescence imaging with spatial resolution beyond the diffraction limit (super-resolution imaging) can be achieved^9^. STED offers two practical advantages: real-time improvement in spatial resolution without any numerical processing, and temporal resolution comparable to that of a scanning confocal laser microscope used as the base system. Recently, fluorescent dyes capable of efficient fluorescence depletion have been developed, improving the usability of STED super-resolution imaging^10–12^. Moreover, the depth resolution of STED can be significantly improved by controlling the wavefront of the erase beam, making it a promising tool for observing biological specimens with dense three-dimensional structures (3D-STED)^13–15^.

In today’s life science research, fluorescent proteins have become indispensable for fluorescence imaging of biological samples, particularly for observing live tissues and cells, and their application to STED imaging is also highly expected^16^. However, a rational method for selecting fluorescent proteins for STED microscopy has not yet been established. The main reason is that research on the FD characteristics of fluorescent proteins is limited, and compatibility between fluorescent proteins and STED systems relies heavily on empirical observations. Therefore, to assess the utility and practicality of fluorescent proteins in STED, it is necessary to establish a method to evaluate spectroscopic properties related to FD.

In STED super-resolution imaging, it is essential that the STED microscopic system can effectively induce FD for the fluorescent molecules used. For this, the erase beam wavelength needs to lie within the stimulated-emission wavelength range of the fluorescent molecule. It is also important to select fluorescent molecules whose stimulated emission cross-section at the depletion wavelength is significantly larger than the S_n_ (n≥2) ← S_1_ absorption cross-section, because the S_n_ (n≥2) ← S_1_ absorption competes with stimulated emission and reduces its efficiency, and can also promote transitions to other states, such as triplet states via S_1_ or S_n_, leading to long-lived dark states and irreversible photobleaching^17^. In addition, fluorescent molecules with low intersystem crossing efficiency from S_1_ to T_1_ are desirable. This is because fluorescent molecules exhibiting high intersystem crossing efficiency from S_1_ to T_1_ are prone to forming long-lived dark states and undergoing photobleaching, making them unsuitable for STED, where maintaining signal strength is critical^18^. Transient absorption spectroscopy (TAS), a two-color spectroscopy (TCS) method that measures absorption spectra in the S_1_ state of fluorescent molecules^17,19^, is well suited to evaluating such spectroscopic characteristics. Furthermore, it is necessary to verify that FD by stimulated emission can be efficiently induced at the output of the STED system’s erase beam, and that secondary emission does not occur upon erase beam irradiation, because insufficient depletion intensity or secondary emission compromises FD and consequently hinders resolution improvement. To address this issue, fluorescence dip spectroscopy (FDS)^20^, another TCS-based method, provides an effective means to examine FD under conditions that closely replicate practical STED observation conditions.

In this study, we assessed the compatibility of widely used red fluorescent proteins (TagRFP, DsRed2, mScarlet) and green fluorescent proteins (EGFP, superfolderGFP, mNeonGreen) with the STED microscope system by evaluating FD-related properties using TAS and FDS. Using the Randomly Interleaved Pulse Train (RIPT) method^21^ as TAS, which can measure the time evolution of transient absorption spectra in the nanosecond region with high precision, we identified the wavelength range of stimulated emission for each fluorescent protein. We also confirmed that, for all fluorescent proteins, stimulated emission is dominant over S_n_ (n≥2) ← S_1_ absorption at the wavelength of the STED system’s erase beam, and found no long-lived intermediates generated upon illumination with the pump beam. We further determined the irradiation intensity required to induce FD using FDS. Finally, we performed 3D-STED super-resolution imaging of biological samples expressing each fluorescent protein identified as compatible with the STED system.

## Results

### Clarification of the wavelength range for the erase beam using TAS

In TAS, molecules are illuminated with a probe beam (corresponding to the erase beam in STED) either before or after being excited from S_0_ to S_1_ by a pump beam. The change in the transmittance of the probe beam before and after excitation is measured as the transient absorption change (Δ_Abs_). When excitation from S_0_ to S_1_ occurs by the pump beam, absorption from S_0_ (ground-state absorption) decreases, so Δ_Abs_ becomes negative (bleach signal). If Δ_Abs_ is more negative than the bleach signal at a given wavelength of the probe beam, it indicates that stimulated emission has occurred. Conversely, a positive Δ_Abs_ indicates that S_n_ (n≥2) ← S_1_ absorption has occurred. Accordingly, measuring Δ_Abs_ using TAS delineates the S_n_ (n≥2) ← S_1_ absorption and stimulated emission bands for each fluorescent protein, enabling evaluation of compatibility with the STED erase-beam wavelength. In addition, by tracking the temporal change of the bleach signal, the generation of long-lived intermediates can be evaluated. Because our STED system uses low-energy 10-ns pump pulses and 20-ns erase pulses, it is important to verify that long-lived intermediates, which reduce efficiency of stimulated emission, are hardly produced during the 20-ns erase beam illumination and thereafter.

In this study, we used the recently developed RIPT method^21^ as TAS, because conventional TAS methods have technical limitations that hinder the simultaneous acquisition of the aforementioned information. Specifically, the pump–probe method using picosecond or femtosecond lasers can neither track the complete relaxation of the S₁ state of fluorescent proteins with S_1_ lifetimes of several nanoseconds nor evaluate the existence of long-lived intermediates because the upper limit of the measurement time window is several nanoseconds. Meanwhile, in continuous-wave TAS, accurate evaluation of stimulated emission signals is hindered by fluorescence background that overwhelms the transient absorption signal, and its modest temporal resolution (∼10 ns) also prevents comprehensive observation of the S_1_ relaxation process. The RIPT method is designed to overcome these technical issues; it enables observation over a time range of ∼100 picoseconds to milliseconds and accurate detection of stimulated emission signals by separating fluorescence from the spectrum.

We first characterized transient absorption of three red fluorescent proteins: mScarlet, TagRFP, and DsRed2. Fig.2(a) shows the result of mScarlet. As shown in Fig. 2(a-1), the spectrum goes negative between 600 nm and 750 nm, where ground-state absorption (and the bleach signal) is very small or negligible, indicating the presence of a stimulated emission band. As shown in Fig. 2(a-2), around 670 nm, the erase beam wavelength of the STED system for red fluorescent proteins (indicated by the red dashed line), Δ_Abs_ is strongly negative, indicating that stimulated emission significantly dominates over competing S_n_ (n≥2) ← S_1_ absorption. Consequently, 670 nm is validated as an appropriate erase beam wavelength for inducing FD in mScarlet. Furthermore, the temporal trace of the transient absorption signal starting at pump illumination in Fig. 2(a-3) shows a decay with lifetime of ∼3 ns, with most of the stimulated emission occurring within ∼15 ns from the rise to the fall. Therefore, the pulse width (20 ns) of the erase beam of the STED system can sufficiently deplete the generated S_1_ state of mScarlet. Additionally, the bleach signal returned to baseline by 15 ns, confirming the absence of long-lived intermediates. Similar results were obtained for TagRFP and DsRed2; see Supplementary Figs.S1(a) and S1(b) for details.

**Figure 1.**
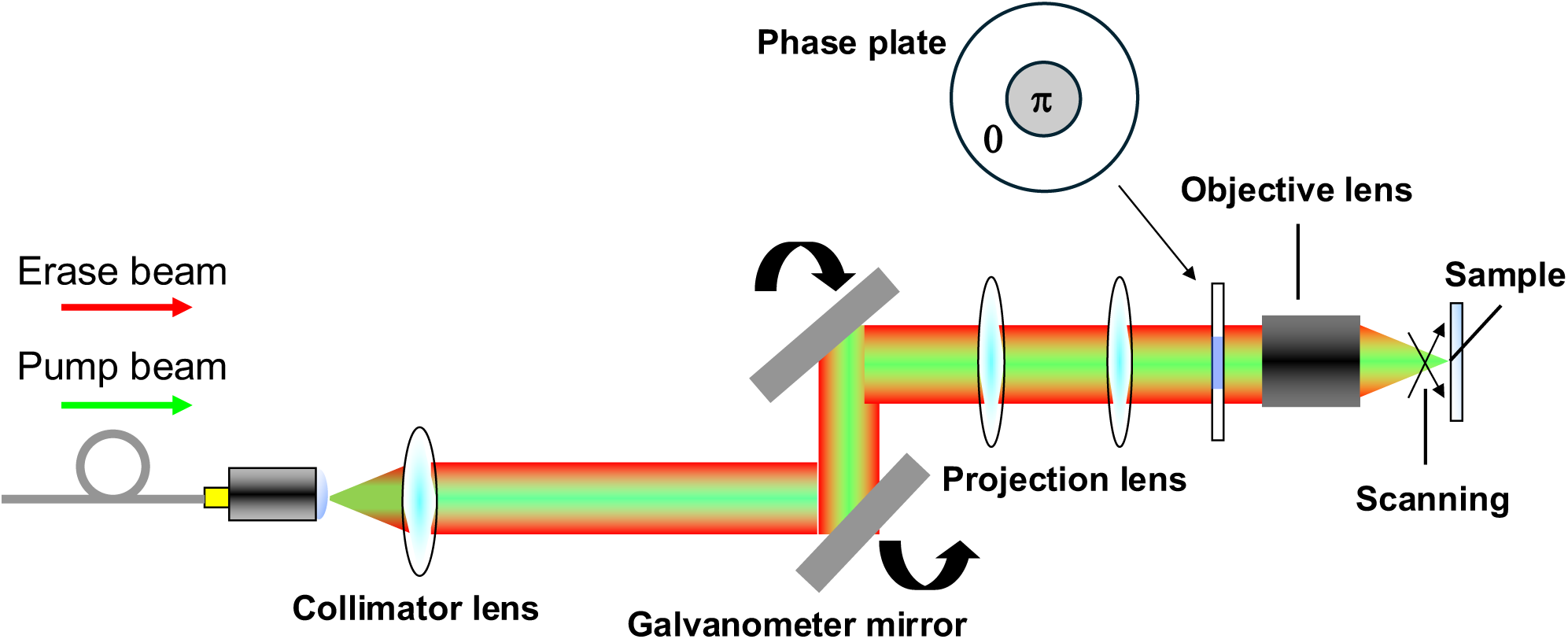
Schematics of 3D-STED super-resolution microscope system.

**Figure 2.**
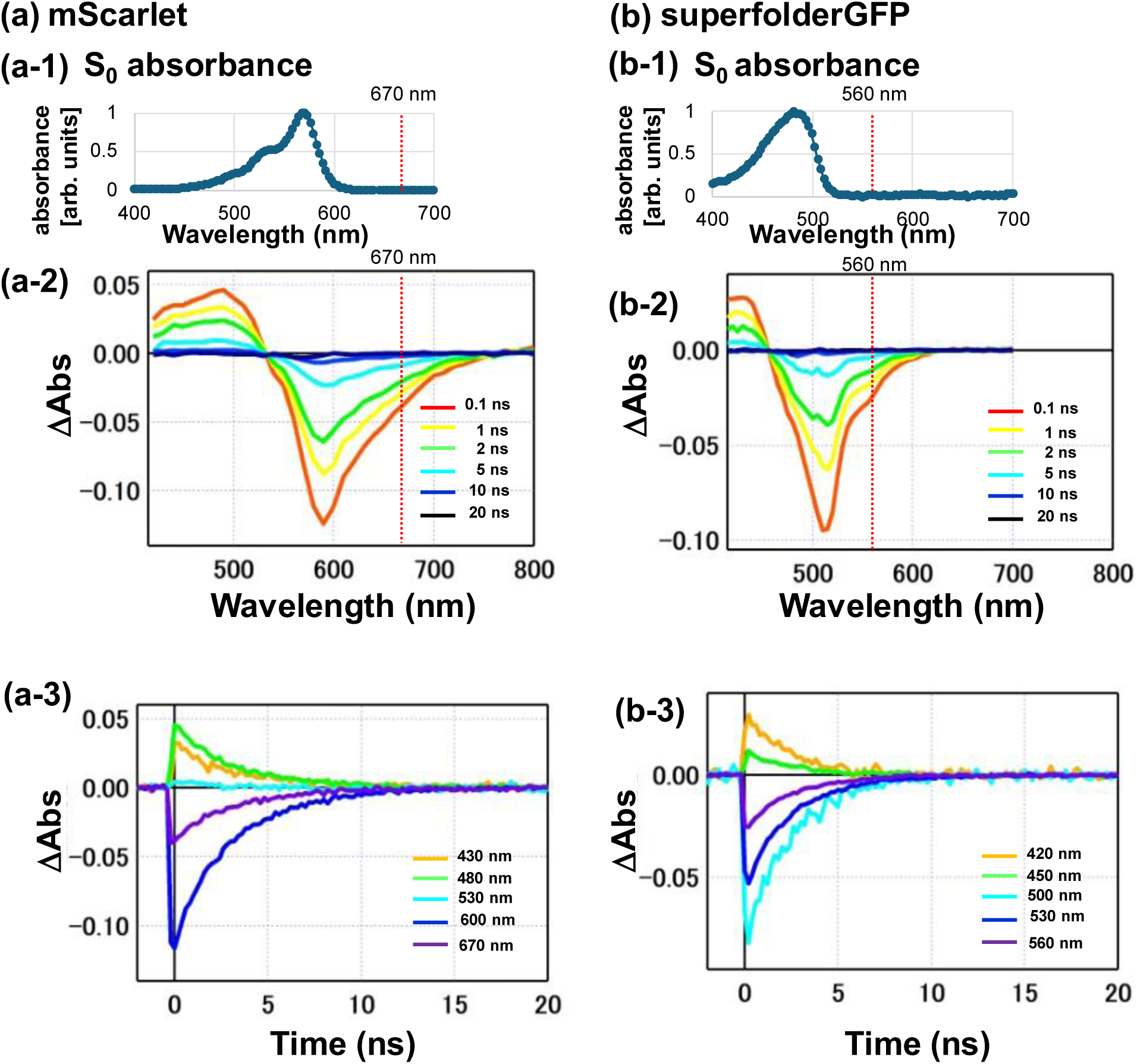
Measured transient absorption spectra of fluorescent protein using the RIPT method. (a) RIPT measurement of mScarlet. (a-1) S_0_ absorption spectrum. The wavelength of the erase beam (670 nm) is indicated by the dashed line. (a-2) Transient absorption spectra at 0.1 ns (red), 1 ns (yellow), 2 ns (light green), 5 ns (light blue), 10 ns (blue), and 20 ns (dark blue) with respect to the time of pump beam irradiation. The wavelength of the erase beam (670 nm) is indicated by the dashed line. (a-3) Time evolution of transient absorption at the wavelengths of 430 nm (orange), 480 nm (green), 530 nm (light blue), 600 nm (blue), and 670 nm (purple). (b) RIPT measurement of superfolderGFP. (b-1) S_0_ absorption spectrum. The wavelength of the erase beam (560 nm) is indicated by a dashed line. (b-2) Transient absorption spectrum at 0.1 ns (red), 1 ns (yellow), 2 ns (light green), 5 ns (light blue), 10 ns (blue), and 20 ns (dark blue) with respect to the time of pump beam irradiation. The wavelength of the erase beam (560 nm) is indicated by a dashed line. (b-3) Time evolution of transient absorption at the wavelengths of 420 nm (orange), 450 nm (green), 500 nm (light blue), 530 nm (blue), and 560 nm (purple).

Next, we investigated the transient absorption of three green fluorescent proteins, superfolderGFP, EGFP, and mNeonGreen. Figure 2(b) shows the results for superfolderGFP. As shown in Fig. 2(b-2), the spectrum goes negative between 530 nm and 620 nm, where ground-state absorption (and the bleach signal) is very small or negligible (Fig.2 (b1)), indicating the presence of a stimulated emission band. The wavelength (560 nm, indicated by the red dashed line) of the STED system’s erase beam for green fluorescent proteins falls within this band. The transient absorption signal around 560 nm is strongly negative, indicating that stimulated emission dominates over S_n_ ← S_1_ absorption. Therefore, 560 nm was confirmed to be an appropriate erase-beam wavelength for inducing FD for superfolderGFP. A lifetime of 3 ns is observed, most of the stimulated emission occurs within 10 ns from the rise to the fall, and the bleach returns to zero by 10 ns (Fig. 2(b-3)), indicating the absence of long-lived intermediates. Similar results were obtained for EGFP and mNeonGreen (Supplementary Figs.S1(c) and S1(d)).

### Evaluation of FD properties of fluorescent proteins

Next, we performed FDS to assess whether FD could be induced efficiently enough to enable 3D-STED super-resolution imaging in each fluorescent protein sample using our STED system. The phase plate was removed from the 3D-STED system, and either a Gaussian pump beam alone or the pump and erase beams were coaxially focused onto each sample (see Methods section for details). Fluorescence intensity of the resulting image was quantified. Fig. 3(a) show the results of FD induction using a sample of fixed LLC-PK1 cells expressing mScarlet-Adhiron6^22^ (also called affimer6^23^) that specifically binds to F-actin. The fluorescence images of the cells scanned with the pump beam alone and scanned with the pump and the erase beams simultaneously are shown in Figs. 3(a-1) and 3(a-2), respectively. The intensity profiles along the dashed lines in Figs. 3(a-1) and 3(a-2) are compared in Fig. 3(a-3). When simultaneously irradiated with the pump and erase beams, the overall fluorescence intensity was significantly reduced, indicating that FD was efficiently induced for mScarlet within the cell, although weak fluorescence emission remained. Similarly, FD was efficiently induced for superfolderGFP fused to F-actin binding affimer6 in PFA-fixed LLC-PK1 cells (Fig.3(b)). The results for other fluorescent proteins are shown in Supplementary Fig.2.

**Figure 3.**
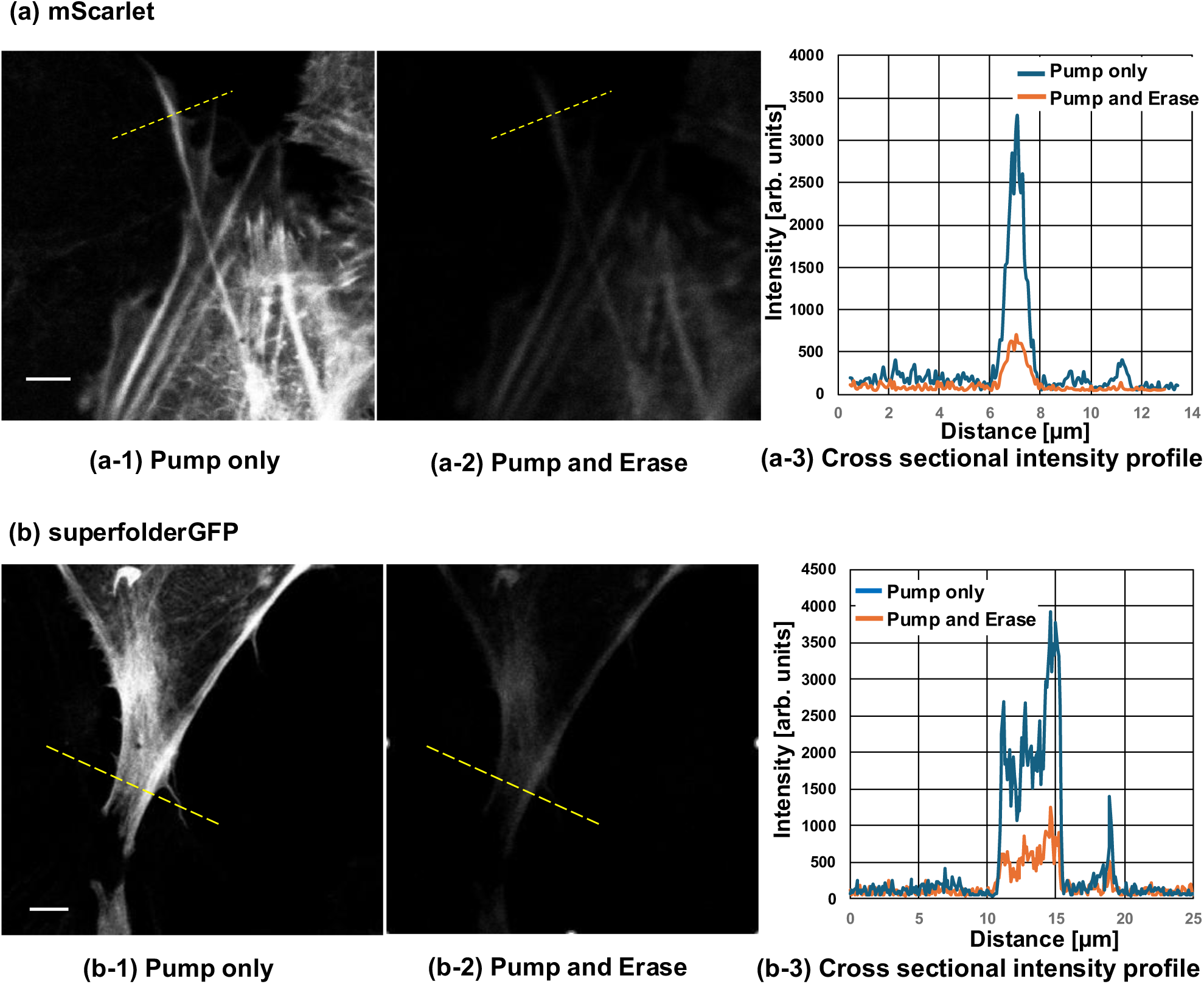
FD induction for mScarlet and superfolderGFP using samples of fixed LLC-PK1 cells. (a) FD induction results for mScarlet. (a-1) Fluorescence image of an LLC-PK1 cell expressing mScarlet fused to affimer6 that specifically binds to F-actin acquired by scanning with the pump beam of the 16 μW peak power alone. Scale bar: 5 μm. (a-2) The fluorescence image of the same field of view as (a-1) acquired by scanning simultaneously with the pump together with the erase beams of the 54 mW peak power. (a-3) Cross-sectional intensity profiles along the dashed lines in (a-1) and (a-2). The blue curve shows the profile for irradiation with the pump beam alone, and the orange curve shows the profiles for simultaneous irradiation with the pump and erase beams. (b) FD induction results for superfolderGFP. (b-1) The fluorescence image of an LLC-PK1 cell expressing superfolderGFP fused to affimer6 that specifically binds to F-actin taken by scanning with the pump beam of the 6 μW peak power alone. (b-2) The fluorescence image of the same view as (b-1) taken by scanning simultaneously with the pump together with the erase beams of the 60 mW peak power. Scale bar: 5 μm. (b-3) Cross-sectional intensity profiles along the dashed lines in (b-1) and (b-2). The blue curve shows the profile for irradiation with the pump beam alone, and the orange curve shows the profiles for simultaneous irradiation with the pump and erase beams.

FD properties of each fluorescent protein were further assessed by plotting the fluorescence intensity as a function of the erase beam intensity introduced into the objective lens. FD properties of the red fluorescent proteins (TagRFP, DsRed2, mScarlet) and the green fluorescent proteins (EGFP, superfolderGFP, and mNeonGreen) were shown in Fig. 4(a) and (b), respectively. For comparison, FD properties of Nile Red and BODIPY in polystyrene beads were also included in Fig. 4(a) and (b), respectively. The results indicate that FD induction occurs for all fluorescent proteins, depending on the intensity of the erase beam, although the reduction in fluorescence intensity was smaller than that observed for Nile Red and BODIPY. Efficient FD was observed for mScarlet, DsRed2, superfolderGFP, and EGFP, with fluorescence intensity reductions exceeding 70% using an erase beam of ∼20 mW. At intensities above 50 mW, reductions of 80% or more were observed for five fluorescent proteins, except for mNeonGreen.

**Figure 4.**
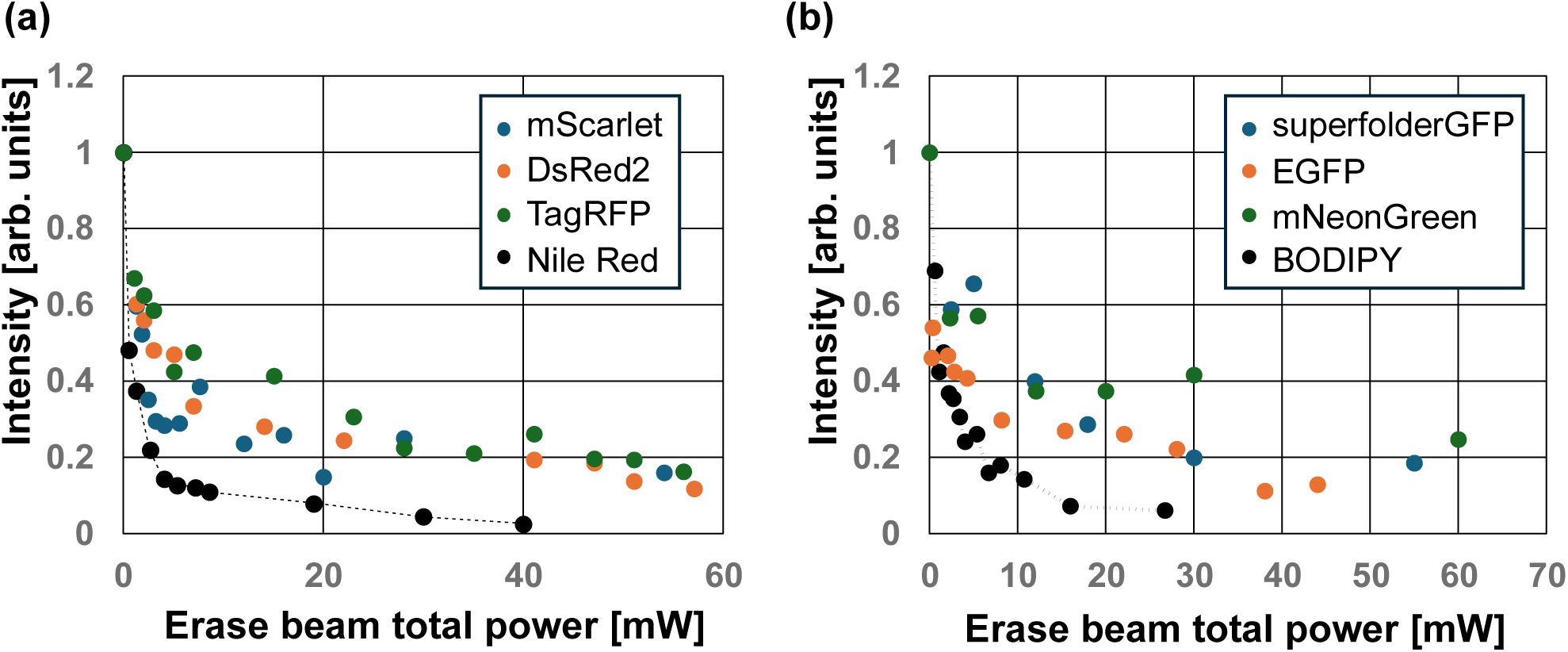
FD properties of the red and green fluorescent proteins. (a) FD plots for TagRFP (green), DsRed2 (orange), mScarlet (blue) and Nile Red (black). As a reference, the Nile Red plot was indicated with the dashed line. (b) FD plots for EGFP (orange), superfolderGFP (blue), mNeonGreen (green) and BODIPY (black). As a reference, the BODIPY plot was indicated with the dashed line.

### Performance assessment of 3D-STED super-resolution system

To assess spatial resolution in our 3D-STED system, we observed three-dimensional fluorescence spots of isolated 60-nm-diameter polystyrene beads containing Nile Red (Fig. 5(a)) with the phase plate inserted into our 3D-STED system. The point spread function (PSF) was measured from these imaged spots. When the beads were irradiated with the pump beam (wavelength: 515 nm, peak power: 14 μW) alone i.e. confocal imaging, the ellipsoidal fluorescent spots with FWHMs of 270 nm in the focal plane and 700 nm in the depth were observed. Conversely, spherical fluorescent spots with a FWHM of 90 nm were obtained through super-resolution imaging by irradiating the beads with the pump beam together with the erase beam (wavelength: 670 nm, peak power: 22 mW). We also observed 75-nm-diameter beads containing BODIPY (Fig. 5(b)). Spherical fluorescent spots with an FWHM of 90 nm were observed through super-resolution imaging by irradiating with the pump beam (wavelength: 473 nm, peak power: 16 μW) and the erase beam (wavelength: 560 nm, peak power: 60 mW). On the other hand, when only the pump beam was irradiated onto the beads, ellipsoidal fluorescent spots with an FWHM of 250 nm at the focal plane and 600 nm in depth were obtained. These results demonstrated that our 3D-STED system provides three-dimensional spatial resolution better than 90 nm.

**Figure 5.**
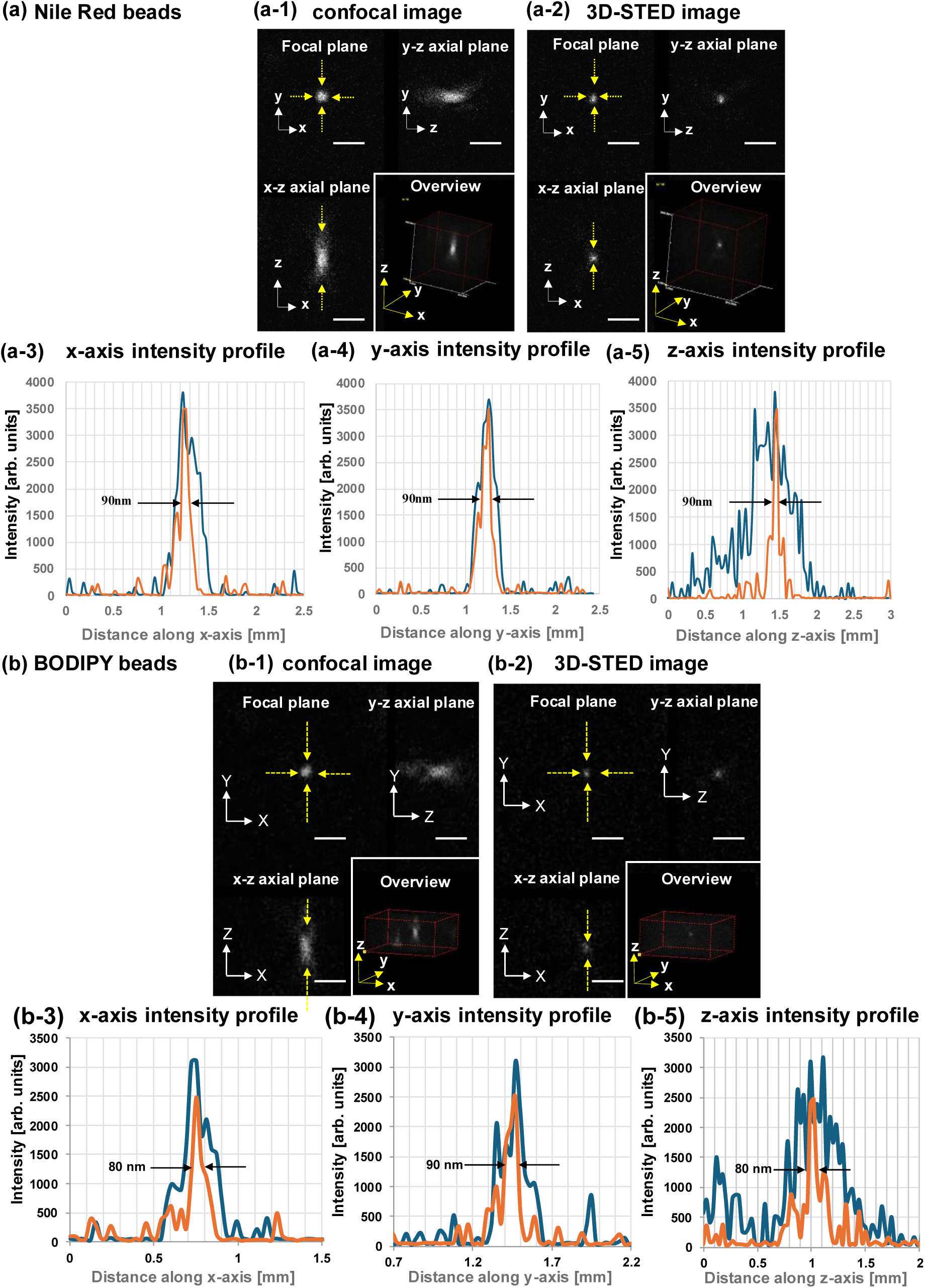
Measured point spread functions in the 3D-STED system by observing isolated fluorescent beads containing Nile Red and BODIPY. (a) PSF measurement results of Nile Red beads. (a-1) and (a-2): Three-dimensional expansion images of the isolated 60-nm-diameter fluorescent beads containing Nile Red irradiated by the pump beam alone (confocal imaging, (a-1)) and simultaneously irradiated the pump and erase beams (3D-STED imaging, (a-2)). Pump beam peak power: 14 μW, erase beam peak power: 22 mW. Scale bar: 1 μm. (a-3), (a-4) and (a-5): The cross-sectional intensity profiles for the beads containing Nile Red along the x-axis in the focal plane (a-3), along the y-axis in the focal plane (a-4), and along the z-axis in the optical axial plane (a-5). (b) PSF measurement results of BODIPY beads. (b-1) and (b-2): Three-dimensional expansion images of the 75-nm-diameter fluorescent beads containing BODIPY in confocal imaging (b-1) and 3D-STED imaging (b-2). Pump beam peak power: 16 μW, erase beam peak power: 60 mW. Scale bar: 1 μm. (b-3), (b-4), (b-5): The cross-sectional intensity profiles for the beads containing BODIPY along the x-axis in the focal plane (b-3), along the y-axis in the focal plane (b-4), and along the z-axis in the optical axial plane (b-5). Blue and orange curves indicate the profiles in confocal and super-resolution imaging, respectively.

To verify the applicability of the 3D-STED system to fluorescent proteins, we similarly observed isolated 100-nm-diameter polystyrene beads coated with purified DsRed2 protein (Fig. 6). Through super-resolution imaging (Fig.6(b)), the fluorescent spots with the FWHM of 150 nm in the focal plane and 100 nm in the depth were obtained. On the other hand, through confocal imaging by irradiating the beads with the pump beam only (Fig.6(a)), the spots with the FWHM of 280 nm in the focal plane and 700 nm in the depth were obtained. The results confirm that the 3D-STED system can provide the three-dimensional resolution surpassing the diffraction limit for fluorescent proteins.

**Figure 6.**
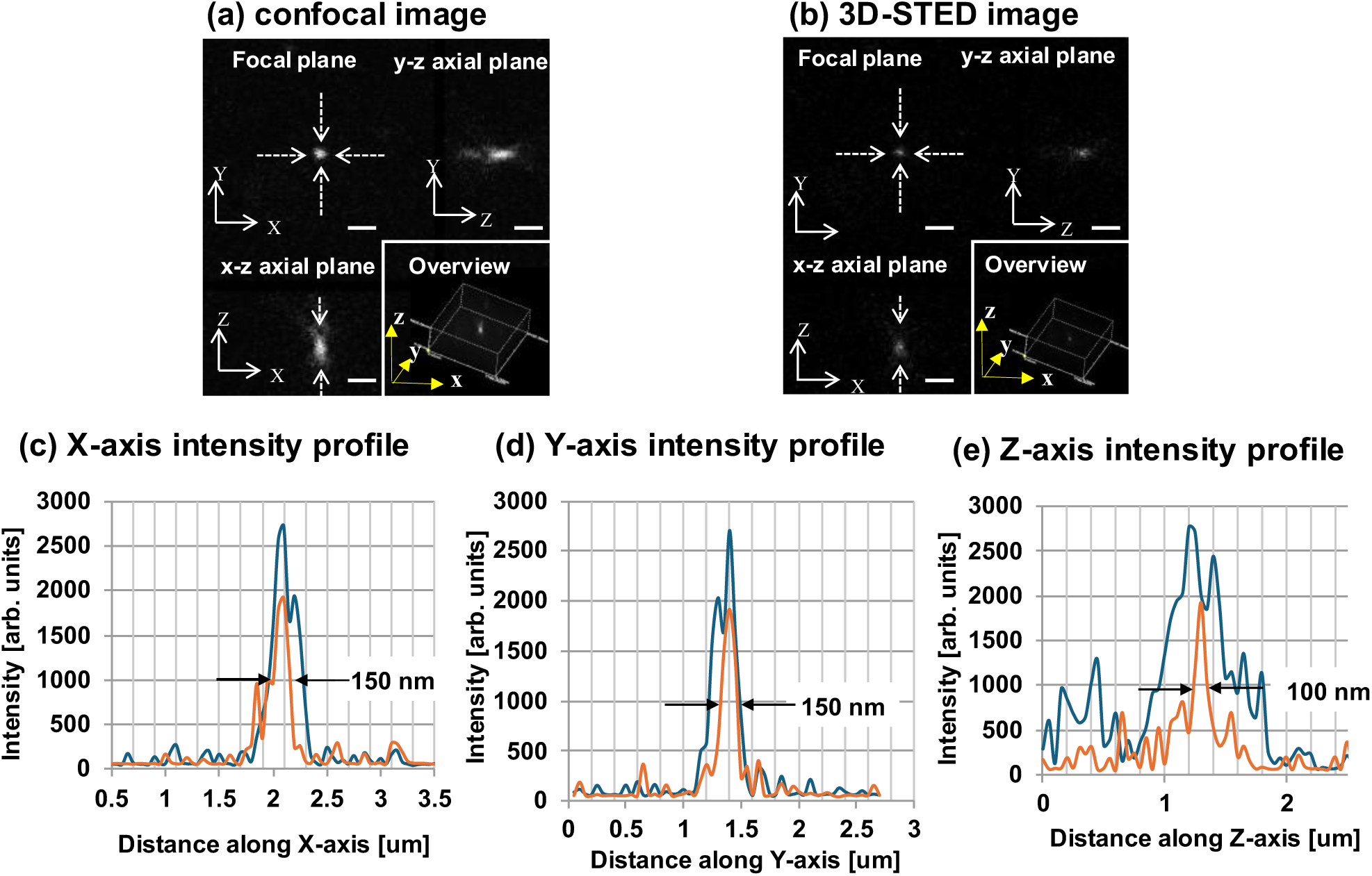
Measured point spread functions in the 3D-STED system by observing 100-nm-diameter polystyrene beads coated with purified DsRed2 protein. (a) and (b): Three-dimensional expansion images of the beads observed in confocal imaging (a) and 3D-STED imaging (b). Pump beam peak power: 3 μW, erase beam peak power: 60 mW. Scale bar: 1 μm. (c) , (d) and (e): The cross-sectional intensity profiles for the beads along the x-axial in the focal plane, along the y-axis in the focal plane, along the z-axis in the optical axial plane, respectively. Blue and orange curves indicate the profiles in confocal and super-resolution imaging, respectively.

### 3D-STED super-resolution imaging of biological samples

Based on the above validation, we selected mScarlet, DsRed2, TagRFP, superfolderGFP, and EGFP as compatible for efficient FD with our 3D-STED system. Biological specimens expressing these five fluorescent proteins were imaged under the illumination conditions shown in Table 1, except for the pump and erase beam intensities. For the red fluorescent proteins, three PFA-fixed samples were prepared: LLC-PK1 cells expressing mScarlet-affimer6 that specifically labels F-actin, rat neurons expressing mitochondria-targeted TagRFP, and rat neurons expressing mitochondria-targeted DsRed2. For the green fluorescent proteins, fixed LLC-PK1 cells expressing superfolderGFP-affimer6 and fixed rat neurons expressing EGFP were prepared. Fig. 7 shows images of cells expressing mScarlet (a), DsRed2 (b), and TagRFP (c). The 3D bird’s-eye view of each image is shown together with a projection view from the direction indicated by arrows ① (xz plane from y-axis), ② (yz plane from x-axis), and ③ (xy plane from z-axis). Fig.8 similarly shows the images of the cells expressing superfolderGFP (a) and EGFP (b). Examination of 3D bird’s-eye view images revealed that the actin filaments and axons imaged with the 3D-STED system were sharper than those imaged with confocal microscopy. The morphology of actin filaments in LLC-PK1 cells or axons of rat neurons was clearly resolved in the projection views of the 3D-STED images, particularly from the x- or y-axis direction (compare ① (xz plane from y-axis) and ② (yz plane from x-axis) views of confocal and 3D-STED images), indicating that the depth resolution was substantially improved.

**Figure 7.**
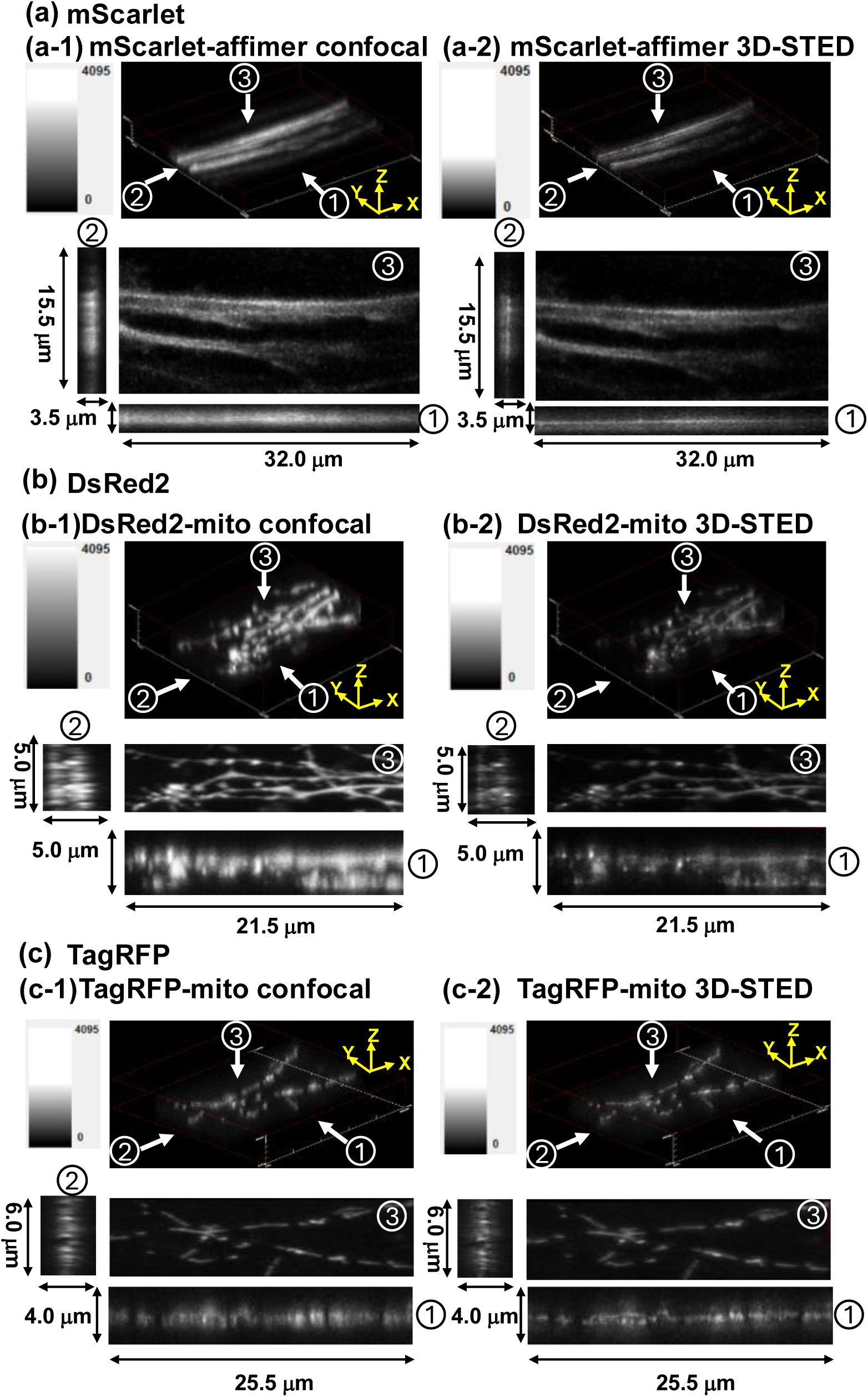
3D-STED super-resolution imaging of cells expressing red fluorescent proteins. (a) Comparison of confocal and 3D-STED images of mScarlet-affimer6 expressing LLC-PK1 cells. (a-1) and (a-2): Three-dimensional images of LLC PK1 cells expressing mScarlet-affimer6 that specifically labels F-actin in confocal imaging (a-1) and 3D-STED imaging (a-2). Pump beam peak power: 16μW, erase beam peak power: 28 mW. The 3D bird’s-eye view is shown together with projection views from the direction indicated by arrows ① (xz plane from y-axis), ② (yz plane from x-axis) and ③ (xy plane from z-axis). (b) Comparison of confocal and 3D-STED images of DsRed2-mito expressing rat neurons. (b-1) and (b-2): Three-dimensional images of rat neurons expressing DsRed2 in confocal imaging (b-1) and 3D-STED imaging (b-2). Pump beam peak power: 12 μW, erase beam peak power: 38 mW. The 3D bird’s-eye view is shown together with projection views from the direction indicated by arrows ① (xz plane from y-axis), ② (yz plane from x-axis) and ③ (xy plane from z-axis). (c) Comparison of confocal and 3D-STED images of TagRFP-mito expressing rat neurons. (c-1) and (c-2): Three-dimensional images of rat neurons expressing mitochondria-targeted TagRFP in confocal imaging (c-1) and 3D-STED imaging (c-2). Pump beam peak power: 16 μW, erase beam peak power: 56 mW. The 3D bird’s-eye view is shown together with projection views from the direction indicated by arrows ① (xz plane from y-axis), ② (yz plane from x-axis) and ③ (xy plane from z-axis).

**Figure 8.**
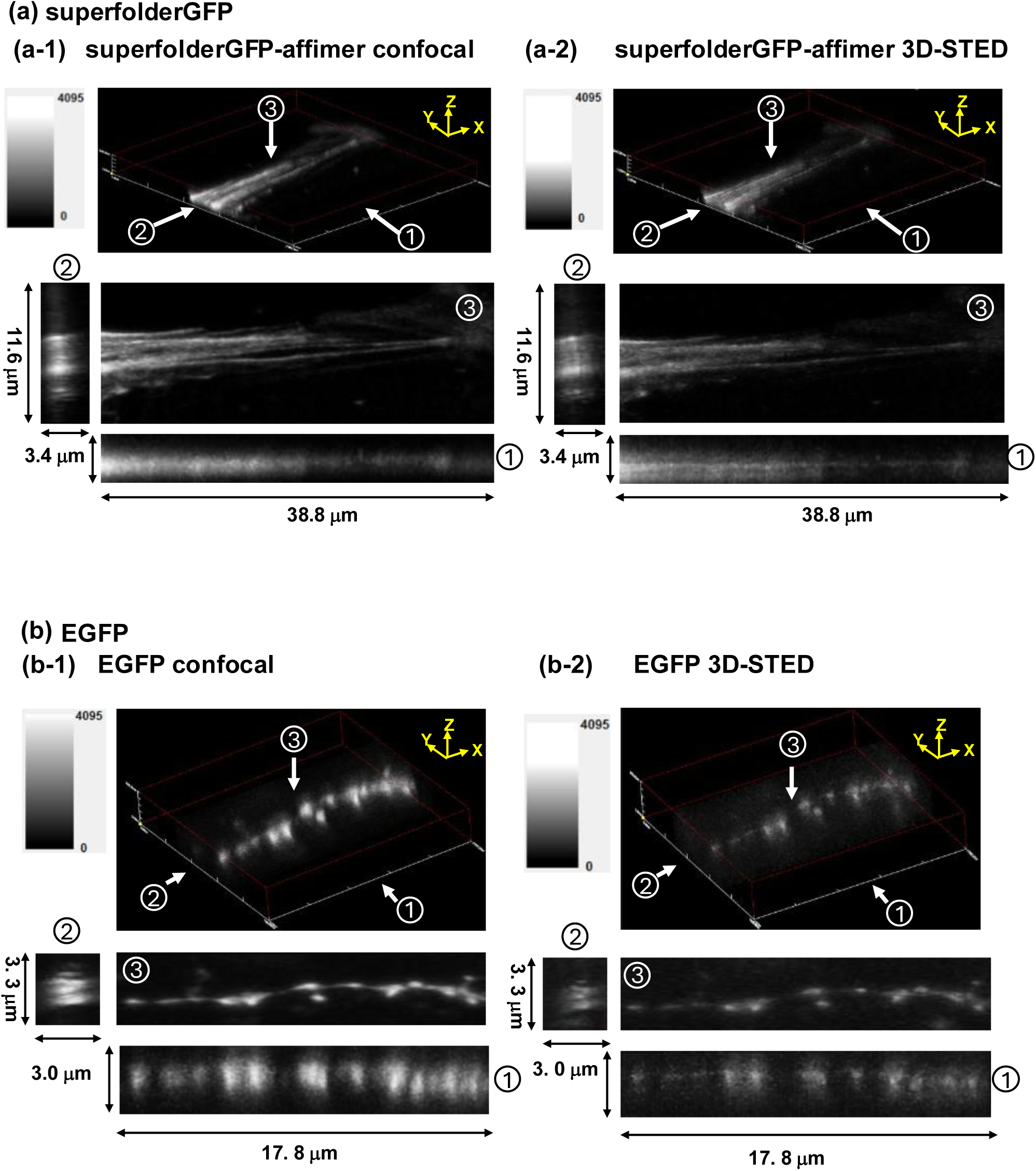
3D-STED super-resolution imaging of cells expressing green fluorescent proteins. (a) Comparison of confocal and 3D-STED images of superfolderGFP-affimer6 expressing LLC-PK1 cells. (a-1) and (a-2): Three-dimensional images of LLC PK1 cells expressing superfolderGFP-affimer6 that specifically labels F-actin in confocal imaging (a-1) and 3D-STED imaging (a-2). Pump beam peak power: 8 μW, erase beam peak power: 48 mW. The 3D bird’s-eye view is shown together with projection views from the direction indicated by arrows ① (xz plane from y-axis), ② (yz plane from x-axis) and ③ (xy plane from z-axis). (b) Comparison of confocal and 3D-STED images of EGFP expressing rat neurons. (b-1) and (b-2) show three-dimensional images of rat neurons expressing EGFP in confocal imaging (b-1) and 3D-STED imaging (b-2). Pump beam peak power: 7μW, erase beam peak power: 50 mW. The 3D bird’s-eye view is shown together with projection views from the direction indicated by arrows ① (xz plane from y-axis), ② (yz plane from x-axis) and ③ (xy plane from z-axis).

**Table 1.**
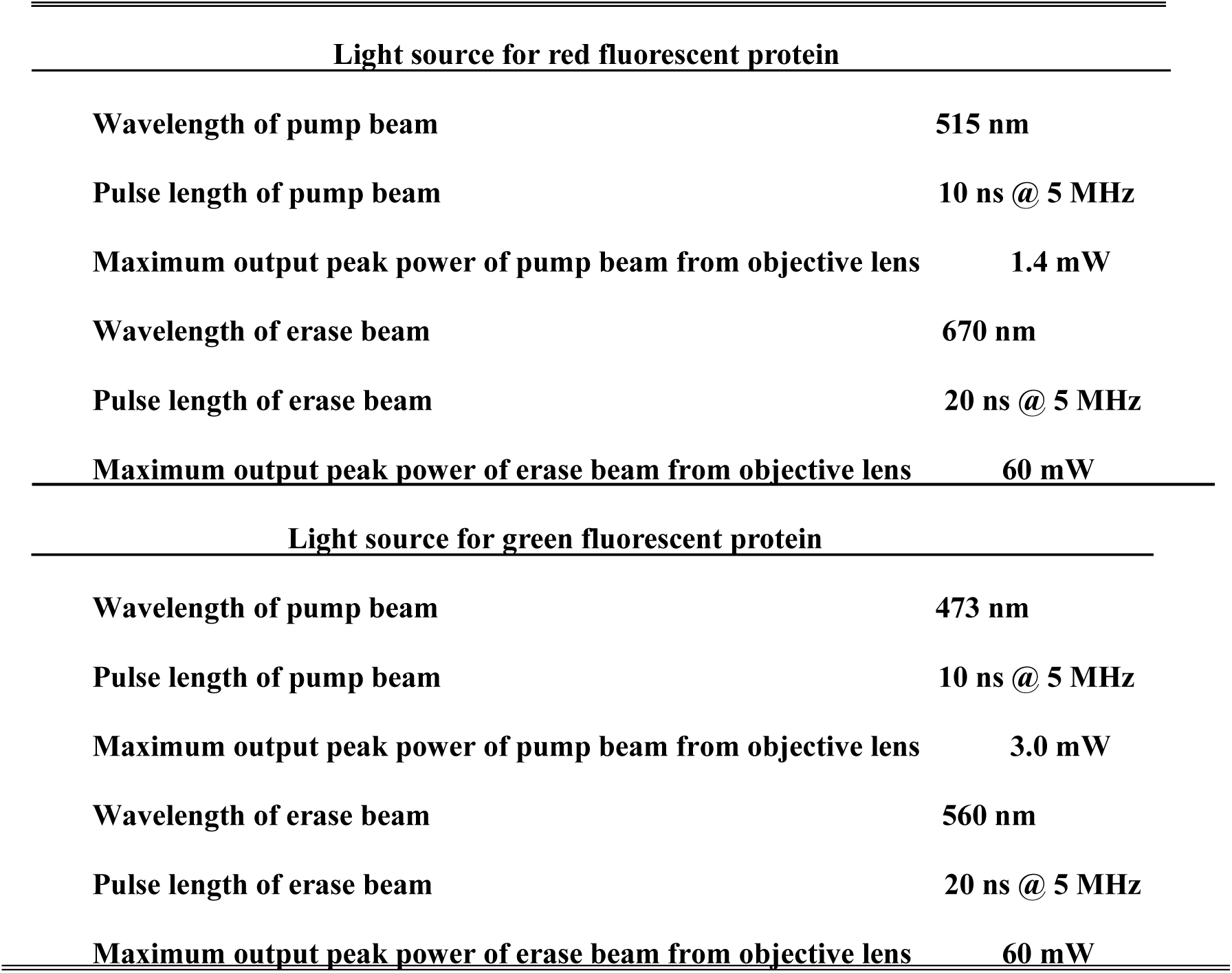
Light source for red fluorescent protein

|  |  |
| --- | --- |
| **Wavelength of pump beam** | **515 nm** |
| **Pulse length of pump beam** | **10 ns @ 5 MHz** |
| **Maximum output peak power of pump beam from objective lens** | **1.4 mW** |
| **Wavelength of erase beam** | **670 nm** |
| **Pulse length of erase beam** | **20 ns @ 5 MHz** |
| **Maximum output peak power of erase beam from objective lens** | **60 mW** |
| **Light source for green fluorescent protein** |  |
| **Wavelength of pump beam** | **473 nm** |
| **Pulse length of pump beam** | **10 ns @ 5 MHz** |
| **Maximum output peak power of pump beam from objective lens** | **3.0 mW** |
| **Wavelength of erase beam** | **560 nm** |
| **Pulse length of erase beam** | **20 ns @ 5 MHz** |
| **Maximum output peak power of erase beam from objective lens** | **60 mW** |

## Discussion

In this study, we demonstrated that the compatibility of fluorescent proteins with the STED microscopy system can be spectroscopically evaluated using TCS. Specifically, TAS analysis using the RIPT method enables identification of the wavelength range of the erase beam available for a given fluorescent protein in STED and investigation of the presence or absence of long-lived intermediates. Measuring FD properties using FDS can assess whether the 3D-STED system can sufficiently induce FD for 3D-STED super-resolution imaging. Indeed, samples expressing fluorescent proteins identified as compatible were imaged using 3D-STED at three-dimensional resolution exceeding the diffraction limit. Although mNeonGreen satisfies the spectral requirement in TAS, its FD efficiency under practical erase-beam intensities was insufficient for reliable 3D-STED imaging in our system.

According to the STED imaging theory^24^, when FD is induced by a nano-second erase beam pulse of the same intensity, a greater reduction in the fluorescence intensity of the fluorophores leads to higher spatial resolution. As shown in Fig. 4, reduction of fluorescence intensity of each fluorescent protein by FD was smaller than that of Nile Red or BODIPY, and the size of the fluorescent spot of the beads coated with DsRed2 (Fig. 6) was larger than that of the beads coated with Nile Red or BODIPY (Fig. 5). These results indicate that in the STED setup used in this study, the induction of FD in fluorescent proteins was less optimized compared to that in fluorescent dyes. In other words, the resolution in STED imaging of fluorescent proteins can be further improved by optimizing FD-induction conditions. For example, using a wavelength-tunable supercontinuum laser that generates high-intensity pulses as a light source for a 3D-STED system, we can closely match the illumination conditions to the FD properties of the fluorescent proteins. Optimization of illumination conditions for efficient FD, guided by TCS measurements presented in this study, is expected to further enhance spatial resolution for fluorescent protein-based STED imaging. In practice, high-quality STED imaging depends not just on FD characteristics, but also on brightness and photostability. The RIPT method is expected to be a useful tool to evaluate these properties. The RIPT method used in this study can monitor not only stimulated emission but also other S_1_-state photoreactions related to the photostability of fluorescent molecules, making it an ideal tool for evaluating the suitability of fluorescent proteins for STED imaging.

In recent years, novel high-performance fluorescent proteins with high brightness and/or photostability have been continuously introduced^25–31^. The TCS measurements used in this study provide a pre-screening strategy for assessing the compatibility with 3D-STED systems for such novel fluorescent proteins. The methodology used in this study is broadly applicable for evaluating STED compatibility not only of fluorescent proteins but also of fluorescent molecules in general. Fundamental spectroscopic data obtained via TCS measurements are expected to facilitate the development of microscopic systems that optimally exploit the optical properties of fluorescent molecules.

## Conclusions

TCS was used to evaluate the compatibility of six fluorescent proteins (mScarlet, TagRFP, DsRed2, superfolderGFP, EGFP, and mNeonGreen) with a STED microscopy system. 3D-STED super-resolution imaging was subsequently conducted on specimens expressing the five proteins identified as compatible, yielding three-dimensional spatial resolution beyond the diffraction limit across all the samples. These results demonstrate that TCS-based measurement of FD-related properties offers a practical approach to the rational selection of fluorescent proteins for STED microscopy. TCS measurements are also expected to provide spectroscopic data to guide the development of optimized microscopic systems.

## Methods

### 3D-STED Super-resolution Microscope System

The 3D-STED super-resolution microscope system (3D-STED system) was set up using the commercially available confocal scanning laser microscope (Olympus: Fluoview 1200) as the base system (Fig. 1). For samples expressing red fluorescent proteins, the semiconductor laser with the wavelength of 515 nm (Cobolt: MLD-515) and the fiber laser with the wavelength of 670 nm (MPB Communications: 2RU-VFL-P-1500-670-B1R) were used for the pump and erase beam light sources, respectively. For green fluorescent protein-expressing samples, the semiconductor laser with the wavelength of 473 nm (Cobolt: MLD-473) and the fiber laser with the wavelength of 560 nm (MPB Communications: 2RU-VFL-P-1500-560-B1R) were used for the pump and erase beam light sources, respectively. The output beams from these sources were intensity-modulated by electro-optic devices and converted into a pulse train with a minimum time width of 10 ns. These beams were transmitted through the same single-mode fiber and coaxially introduced into the illumination optical system of the Fluoview 1200. Table I shows the illumination conditions of the sources. For super-resolution imaging, an annular phase plate for two wavelengths was attached to the objective lens. The phase plate for observing the red fluorescent protein-expressing samples was made by depositing a 2200-nm-thick SiO_2_ layer (refractive index: 1.46) on the central disk area of a quartz substrate polished with a surface accuracy of λ/12 ^32^. A phase plate with a central disk area deposited with a 3050-nm-thick SiO_2_ layer was used for observing the green fluorescent protein-expressing samples. When the pump and erase beams coaxially pass through these plates, only the phase of the erase beam is inverted at the central disk area, creating a three-dimensional fine dark hole caused by destructive interference, while the phase of the pump beam is unchanged. By attaching the phase plate to the objective lens of the 3D-STED system, the pump beam is focused as a Gaussian beam at the center of the dark hole of the erase beam without any misalignment, achieving the illumination for 3D-STED super-resolution imaging. In the 3D-STED system, we used a Si oil objective lens with NA of 1.3 (magnification: 60x). 3D super-resolution imaging was performed by scanning the pump and erase beams using FV-ASW Ver.04.02, the standard software for FV1200.

### Randomly Interleaved Pulse Train (RIPT) Method

The solution of purified fluorescent protein with an absorbance of 0.7 was filled in a quartz cell with a path length of 2 mm. The pump beam, with a pulse width of 25 ps and a repetition rate of 1 kHz, was generated from the OPO laser. Red and green fluorescent proteins were excited with a pulse of 3 μJ at the wavelengths of 550 nm and 470 nm, respectively. The supercontinuum light source with a pulse width of 50–100 ps and a frequency of 20 MHz was monochromatized and used for the probe beam (equivalent to the erase beam in STED). The pump and probe beams were made to pass coaxially through the quartz cell. Delay times between a pump pulse and multiple probe pulses were calculated from those timing signals acquired with a fast oscilloscope, and transient absorption (Δ_Abs_) at each time was calculated from the transmitted light signal. By iterating this procedure with randomly interleaved delay times, a transient absorption decay curve was constructed. The transient absorption spectra were reconstructed by scanning the wavelength.

### Fluorescence Dip Spectroscopy (FDS)

To perform fluorescence dip spectroscopy (FDS), the two-color annular phase plate was removed from the objective lens of the 3D-STED system. We first irradiated the pump beam onto the fluorescent protein-expressing sample alone, and the fluorescence intensity of the image without FD was measured. Subsequently, the Gaussian-shaped erase beam was focused onto the sample simultaneously with the pump beam. By varying the intensity of the erase beam, the FD characteristics of the fluorescent proteins at the wavelength of the erase beam were obtained^33^.

### Preparation of Recombinant Proteins

The cDNA sequences encoding each fluorescent protein were inserted downstream of the His-tag in the bacterial expression vector pRSET B, and *E. coli* JM109 (DE3) was transformed with these plasmids. For proteins other than mNeonGreen, colonies obtained from the transformation were pre-cultured overnight at 37 ℃ in LB medium, then diluted 1:1000 in LB medium and cultured overnight at 37 ℃. For mNeonGreen, the colonies were directly cultured overnight at 37 ℃. Cultured *E. coli* cells were collected by centrifugation, resuspended in TN buffer (50 mM Tris-HCl, 300 mM NaCl, pH 7.4), and disrupted by sonication. The proteins were purified from cell extracts by affinity purification using Ni Sepharose 6 Fast Flow (Cytiva). Imidazole used for elution was removed using a PD-10 column (Cytiva), and the resulting solution was used as the sample for downstream applications, including RIPT measurements. Absorbance was measured using NanoDrop (ThermoFisher Scientific).

### Cell Culture

HeLa cells (a generous gift from Dr. Yoshimori, Osaka University) and LLC-PK1 cells (EC86121112-F0, ECACC) were cultured in DMEM high glucose (nacalai tesque) containing 10% FBS or Medium 199 (Merck) containing 10% FBS, respectively, at 37°C in a 5% CO_2_ environment.

### Gene Delivery into Cultured Cells

The cDNA sequences encoding cpGFP-affimer6^22^, affimer6-mScarlet, and affimer6-mNeonGreen were inserted into the modified pcDNA3 mammalian expression vector using standard molecular cloning methods. cDNAs of mScarlet^34^ and mNeonGreen^35^ were purchased from Addgene. HeLa cells were transfected using Lipofectamine 3000 (ThermoFisher Scientific), and LLC-PK1 cells were transfected using the NEPA21 electroporator (Nepagene).

### Preparation of Microscopy Specimens from Cultured Cells

Transfected HeLa and LLC-PK1 cells were cultured on collagen-coated (Cellmatrix type I-C, Nitta Gelatin) cover glasses (ThermoFisher Scientific) and cultured for 1 day. Cells were then fixed in PBS containing 3% paraformaldehyde (PFA) at room temperature for 30 min, washed three times with PBS, and mounted on glass slides using ProLong Glass Antifade Mountant (ThermoFisher Scientific).

### Primary Culture of Rat Hippocampal Neurons and Preparation of Microscopy Specimens

Hippocampi were dissected from Wistar rat embryos (E17) and chopped into small pieces. The cell population consisting mainly of neurons and astrocytes was isolated using neuron dissociation solutions S (FUJIFILM Wako Pure Chemical Corporation) following manufacturer’s instruction. Cells (1x10^5^) were seeded into flexiperm Disc (Sarstedt) wells on poly-L-lysine-coated cover glasses and cultured overnight in MEM (Merck) containing 5% horse serum, 5% fetal bovine serum, and 1 µM pyruvate at 37 ℃ in 5% CO_2_ incubator. The medium was then replaced with MEM containing 2 mM L-glutamine, 2% B27 Electro supplement (ThermoFisher Scientific), and 50 µM DL-aminophosphonovalerate, and the culture was continued. On DIV 7, cells were transfected with TagRFP-mito plasmid (Evrogen), DsRed2-mito plasmid (Takara Bio), or pGL3-homer1-EGFP plasmid using Lipofectamine 2000 (ThermoFisher Scientific). On DIV 20, cells were washed with warm PBS and fixed with PBS containing 4% paraformaldehyde (pH 7.4) at room temperature for 30 min. After washing three times with PBS, cells were air-dried and mounted on slide glasses with ProLong Diamond Antifade Mountant (ThermoFisher Scientific) and stored at 4°C.

### Resolution Evaluation of the 3D-STED System Using Fluorescent Beads

Three-dimensional resolution of the 3D-STED super-resolution microscopy system was evaluated using isolated fluorescent beads containing Nile Red (R60: ThermoFisher Scientific) with the average particle size of 60 nm in diameter and beads containing BODIPY (G75: ThermoFisher Scientific) with the average particle size of 75 nm in diameter. Both fluorescent beads are smaller than the diffraction limit. The beads were scattered on a slide glass plate at a particle concentration of 10 particles/μm^2^ and sealed with a cover glass using a silicone adhesive.

## Acknowledgements

Funding: Japan Society for the Promotion of Science (JSPS) Grant-in-Aid for Scientific Research (KAKENHI) (C) and (B) (18K06819, 21K06750 to K.S., 17H04013, 20H03412 to S.T.); JSPS Fund for the Promotion of Joint International Research (Fostering Joint International Research (B)) (18KK0222 to S.T.); We would like to express sincere gratitude to Prof. Mitsuru Jimbo of Kitasato University School of Marine Biosciences for providing guidance on the preparation of fluorescent beads coated with fluorescent proteins;

## Author contributions

K.S. prepared LLC-PK1 cell samples and conducted FDS and STED imaging using these samples. D.O. prepared rat neuron samples and conducted FDS and STED imaging using these samples. A.S. prepared purified fluorescent proteins for RIPT. T.N. conducted RIPT experiments and analyzed the data. H.K. designed experiments in discussion with other collaborators. Y.I. set up the STED system and conducted FDS and STED imaging with K.S. and D.O.,and analyzed the data. S.T. supervised the project. K.S. and Y.I. jointly wrote the manuscript with inputs from all authors.

## Data availability statement

All data are included in the manuscript and supporting information. Raw data for RIPT experiments and microscopic images are available from the corresponding author upon reasonable request.

