## supplementary Fig for "Evaluation of fluorescent proteins for compatibility with STED microscopy systems using two-color spectroscopies"

[5]Present address: Department of Anatomy and Neurobiology, National Defense Medical College, 3-2 Namiki, Tokorozawa, Saitama, 359-8513, Japan

[8]Present address: Faculty of Distribution and Logistics Systems, Ryutsu Keizai University, 3-2-1 Shin-Matsudo, Matsudo-shi, Chiba 270-8555, Japan

*Corresponding authors

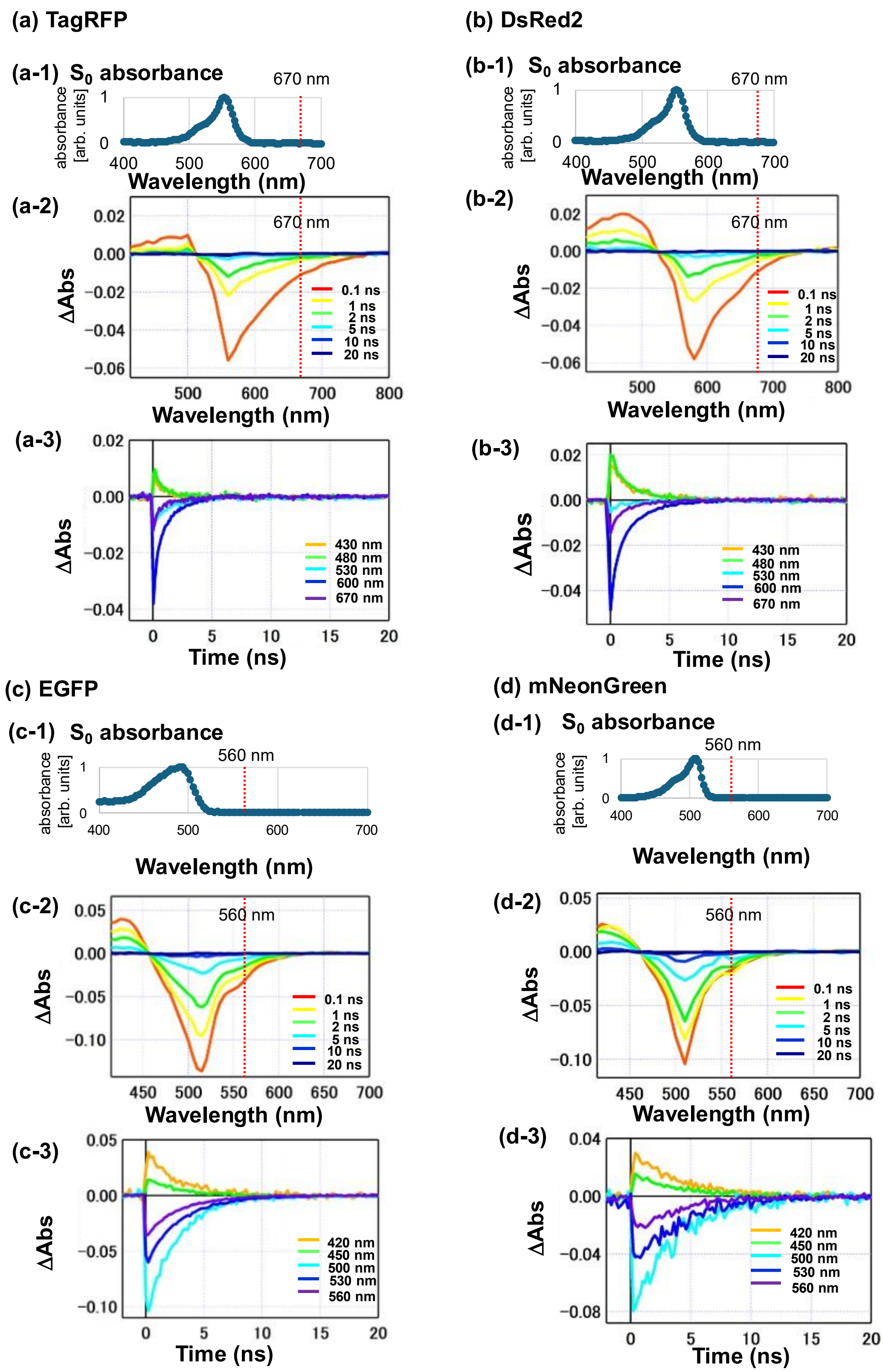


**Supplementary Figure S1. Measured transient absorption spectra of fluorescent protein using RIPT and time evolution of $\Delta_{Abs}$ with respect to the time of pump beam irradiation.**

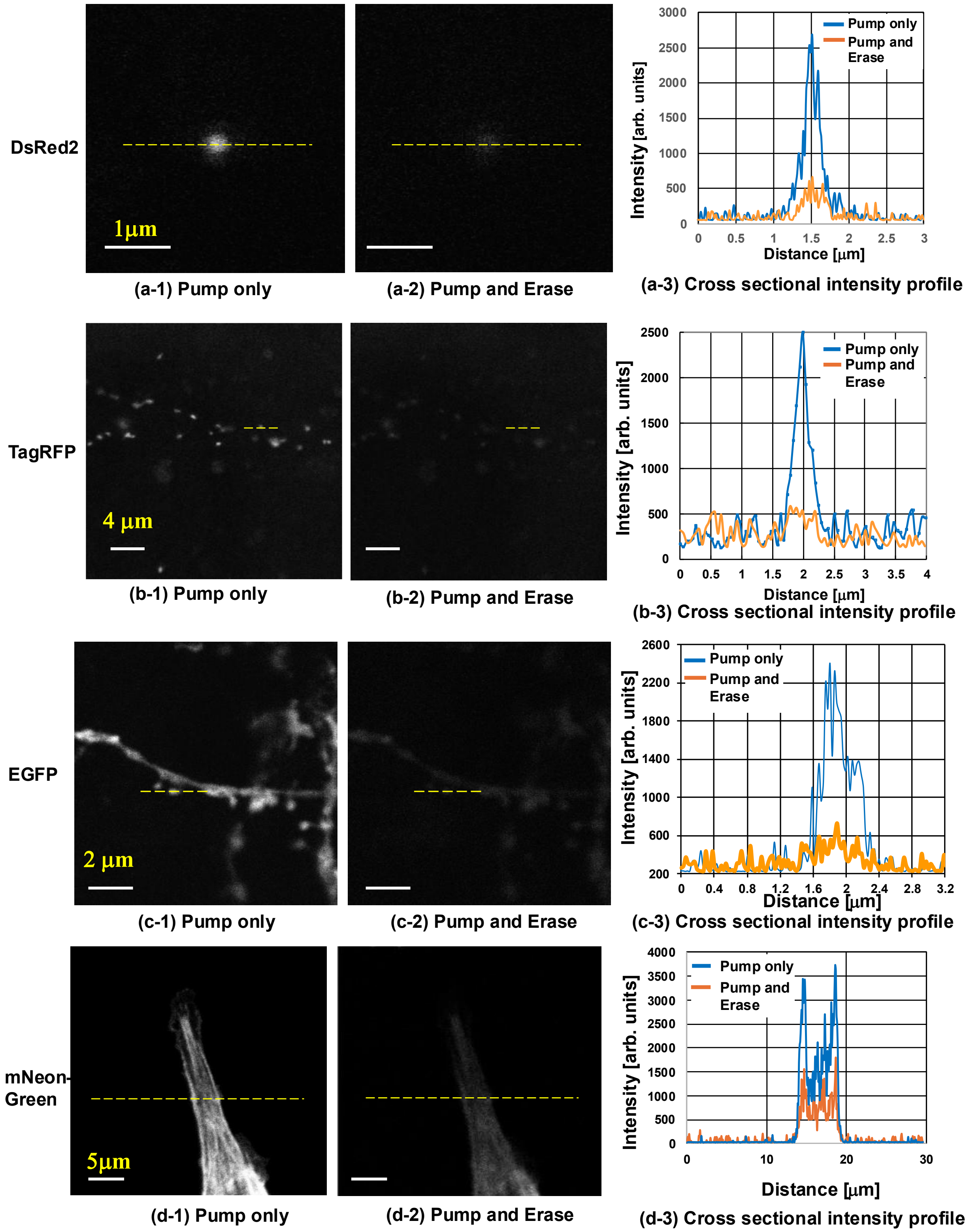


**Supplementary Figure S2. FD induction for fluorescent proteins.**

**Figure Legends**

**Supplementary Figure S1. Measured transient absorption spectra of fluorescent protein using RIPT and time evolution of $\Delta_{Abs}$ with respect to the time of pump beam irradiation.**

(a) Transient absorption measurement of TagRFP.

(a-1) $S_0$ absorption spectrum. The wavelength of the erase beam (670 nm) is indicated by the dashed line.

(a-2) Transient absorption spectrum at 0.1 ns (orange), 1 ns (yellow), 2 ns (light green), 5 ns (light blue), 10 ns (blue), and 20 ns (dark blue) with respect to the time of pump beam irradiation.

(a-3) Time evolution of transient absorption at the wavelengths of 430 nm (orange), 480 nm (green), 530 nm (light blue), 600 nm (blue), and 670 nm (purple).

(b) Transient absorption measurement of DsRed2.

(b-1) $S_0$ absorption spectrum. The wavelength of the erase beam (670 nm) is indicated by the dashed line.

(b-2) Transient absorption spectrum at 0.1 ns (orange), 1 ns (yellow), 2 ns (light green), 5 ns (light blue), 10 ns (blue), and 20 ns (dark blue) with respect to the time of pump beam irradiation.

(b-3) Time evolution of transient absorption at the wavelengths of 430 nm, 480 nm, 530 nm, 600 nm, and 670 nm.

(c) Transient absorption measurement of EGFP.

(c-1) $S_0$ absorption spectrum. The wavelength of the erase beam (560 nm) is indicated by a dashed line.

(c-2) Transient absorption spectrum at 0.1 ns (orange), 1 ns (yellow), 2 ns (light green), 5 ns (light blue), 10 ns (blue), and 20 ns (dark blue) with respect to the time of pump beam irradiation.

(c-3) Time evolution of transient absorption at the wavelengths of 420 nm (orange), 450 nm (green), 500 nm (light blue), 530 nm (blue), and 560 nm (purple).

(d) Transient absorption measurement of mNeonGreen.

(d-1) $S_0$ absorption spectrum. The wavelength of the erase beam (560 nm) is indicated by a dashed line.

(d-2) Transient absorption spectrum at 0.1 ns (orange), 1 ns (yellow), 2 ns (light green), 5 ns (light blue), 10 ns (blue), and 20 ns (dark blue) with respect to the time of pump beam irradiation.

(d-3) Time evolution of transient absorption at the wavelengths of 420 nm (orange), 450 nm (green), 500 nm (light blue), 530 nm (blue), and 560 nm (purple).

**Supplementary Figure S2. FD induction for fluorescent proteins.**

(a-1) A 100-nm-diameter polystyrene bead coated with purified DsRed2 protein was scanned with the pump beam alone, with the peak power of 16 μW.

(a-2) The same DsRed2-coated bead imaged in (a-1) was scanned with the pump beam with the peak power of 16 μW and the erase beam with the peak power of 37 mW.

(a-3) Cross-sectional intensity profiles along the dashed line in (a-1, blue curve) and (a-2, orange curve).

(b-1) A rat neuron expressing mitochondria-targeted TagRFP was scanned with the pump beam alone, with the peak power of 5.2 μW.

(b-2) The same rat neuron imaged in (b-1) was scanned with the pump beam with the peak power of 5.2 μW and the erase beam with the peak power of 28 mW.

(b-3) Cross-sectional intensity profiles along the dashed line in (b-1, blue curve) and (b-2, orange curve).

(c-1) An LLC-PK1 cell expressing mNeonGreen fused to F-actin binding affimer6 was scanned with the pump beam alone, with the peak power of 8 μW.

(c-2) The same LLC-PK1 cell imaged in (c-1) was scanned with the pump beam with the peak power of 8 μW and the erase beam with the peak power of 60 mW.

(c-3) Cross-sectional intensity profiles along the dashed line in (c-1, blue curve) and (c-2, orange curve).

(d-1) A rat neuron expressing EGFP was scanned with the pump beam alone, with the peak power of 12 μW.

(d-2) The same rat neuron imaged in (d-1) was scanned with the pump beam with the peak power of 12 μW and the erase beam with the peak power of 15.4 mW.

(d-3) Cross-sectional intensity profiles along the dashed line in (d-1, blue curve) and (d-2, orange curve).
